# Prebiotics and community composition influence gas production of the human gut microbiota

**DOI:** 10.1101/2020.01.30.928085

**Authors:** Xiaoqian Yu, Thomas Gurry, Le Thanh Tu Nguyen, Hunter S. Richardson, Eric J. Alm

## Abstract

Prebiotics confer benefits to human health often by promoting the growth of gut bacteria that produce metabolites valuable to the human body, such as short chain fatty acids (SCFAs). While prebiotic selection has strongly focused on maximizing the production of SCFAs, less attention has been paid to gases, a byproduct of SCFA production that also has physiological effects on the human body. Here, we investigate how the content and volume of gas production by human gut microbiota is affected by the chemical composition of the prebiotic and by the composition of the microbiota. We first constructed a linear systems model based on mass and electron balance and compared the theoretical product range of two prebiotics, inulin and pectin. Modeling shows that pectin is more restricted in product space, with less potential for H_2_ but more potential for CO_2_ production. An *ex vivo* experimental system showed pectin degradation produced significantly less H_2_ than inulin, but CO_2_ production fell outside the theoretical product range, suggesting fermentation of fecal debris. Microbial community composition also impacted results: methane production was dependent on the presence of *Methanobacteria*, while inter-individual differences in H_2_ production during inulin degradation was driven by a *Lachnospiraceae* taxon. Overall, these results suggest that both the chemistry of the prebiotic and the composition of the microbiota are relevant to gas production. Metabolic processes that are relatively prevalent in the microbiome, such as H_2_ production will depend more on substrate, while rare metabolisms like methanogenesis depend more strongly on microbiome composition.

**Importance:** Prebiotic fermentation in the gut often leads to the co-production of short chain fatty acids (SCFAs) and gases. While excess gas production can be a potential problem for those with functional gut disorders, gas production is rarely taken into account during prebiotic design. In this study, we combined the use of theoretical models and an *ex vivo* experimental platform to illustrate that both the chemical composition of the prebiotic and the community composition of the human gut microbiota can affect the volume and content of gas production during prebiotic fermentation. Specifically, more prevalent metabolic processes such as hydrogen production was strongly affected by the oxidation state of the probiotic, while rare metabolisms such as methane production was less affected by the chemical nature of the substrate and entirely dependent on the presence of *Methanobacteria* in the microbiota.

## Introduction

The gut microbiota plays an important role in human nutrition and health, leading to increasing interest in modulation of the gut microbiome via dietary interventions for improving human health (1–3). Compounds that can be selectively metabolized by microbes in the gut resulting in beneficial effects on the host are defined as prebiotics (4). While some phenolic compounds and fatty acids are suspected to have prebiotic activities, most known prebiotics are dietary carbohydrates that are neither digested or absorbed in the human small intestine, thus capable of reaching the colon and promoting the growth of selective beneficial bacteria (4, 5). These bacteria, in turn, can prevent the colonization of pathogens or produce metabolites that are beneficial for the human body, mostly notably short chain fatty acids (SCFAs) such as acetate, propionate and butyrate. These SCFAs not only contribute directly to host energy metabolism but have a number of positive effects on host physiology. Butyrate is the major energy source for colonocytes and enterocytes (6), and can also activate gluconeogenesis, modulate inflammatory responses and cytokine levels via G protein-coupled receptors or histone deacetylases (7). Similarly, acetate and propionate are involved in the regulation of host immune or metabolic systems (7, 8). Thus, selection for prebiotics has largely focused on those that allow the proliferation of bacteria that maximize production of SCFAs (5, 9, 10).

SCFA fermentation from carbohydrates by the gut microbiota is often coupled with the production of gases. Production of H_2_ is often necessary for the cycling of NAD^+^/NADH during fermentation, and CO_2_ is released whenever decarboxylation occurs (11). H_2_ can be further utilized by methanogens and sulfate reducers for the production of CH_4_ and H_2_S (12). Most intestinal gas is absorbed into the bloodstream and removed *via* the lungs (13), but it can still have physiological effects on the human body. The volume of gas production can affect the distension of the colonic wall and in turn affect the speed of material transition through the colon (14). Methane production can result in slowed intestinal transit and reduced serotonin levels in the gastrointestinal tract, potentially impacting constipation predominant irritable bowel syndrome (IBS-C) and chronic constipation (15). Therefore, gas production may be an important factor to consider in the selection of prebiotics, especially since bloating is a major symptom for many functional gut disorders such as IBS (16).

Many prebiotics are already known to impact fermentation products. For example, short-chain fructooligosacchaides (FOS) and inulin are some of the most extensively documented prebiotics because they promote the growth of Bifidobacteria and increase SCFA production (4, 5). In addition, the low FODMAP (fermentable oligosaccharides, disaccharides, monosaccharides, and polyols) diet that has been shown to improve the symptoms of some IBS patients, because foods containing FOS and inulin can increase luminal distension and gas production (17, 18). Thus, it may be valuable to identify prebiotics that maximize SCFA and minimize gas production, or minimizes the production of specific gases. However, few studies that consider the efficacy of prebiotics simultaneously take gas and SCFA production into account, and systematic investigations on factors that affect gas production in prebiotic fermentation are lacking.

In this study, we investigate whether the chemical composition of the prebiotic and heterogeneity in the composition of gut microbiota can affect the content and volume of gas production during prebiotic fermentation. We compare the fermentation products of two common prebiotics, inulin and pectin, both theoretically *via* linear systems modeling and experimentally *via* an *ex vivo* framework that measures gas and SCFA production of stool microbiota responding to fiber addition (19). We find that inulin, a more reduced carbohydrate, produces more H_2_ compared to pectin, but the amount of H_2_ production is strongly associated with a *Lachnospiraceae* amplicon sequencing variant (ASV). Inulin also yielded greater amounts of the more reduced SCFA butyrate and less acetate. Methane production is, however, less affected by the chemical nature of the substrate, being entirely dependent on the level of *Methanobacteria* in the microbiota. Overall, these results suggest that the production of different gases upon prebiotic fermentation by gut microbiota are differentially affected by the chemical nature of the prebiotic and microbiome composition.

## Results

### Modeling community production with mass and electron balance

To explore the general effect of prebiotic chemical composition on fermentation product formation, we established a linear systems model that allowed us to determine the theoretical range of product output considering mass and electron balance. Considering a system of **n** chemicals as possible inputs and outputs, made up of a total of **m** chemical elements, we defined a matrix **M** in which the rows represent different elements and columns represent different chemicals; the elemental composition of a chemical is thus a column in **M**. The total number of valence electrons in the chemical is also counted as an “element”, and consists of a separate row in **M**. Thus, any reaction that satisfies both mass and electron balance is a n-dimensional vector **s**, whose elements are the stoichiometric coefficients of the chemicals in **M**, and satisfy **Ms=0** (See Figure 1a for a more detailed representation). By definition, **s** must be within the null space of **M**. The feasible product space of the biological system, represented by the elements in **s** that are coefficients of the possible products, is thus a convex cone defined by the linear combinations of the basis vector of null(**M**). Since our model did not account for the thermodynamic constraints on the metabolic fluxes within the system, it represented an upper limit of the feasible product space.

**Figure 1.**
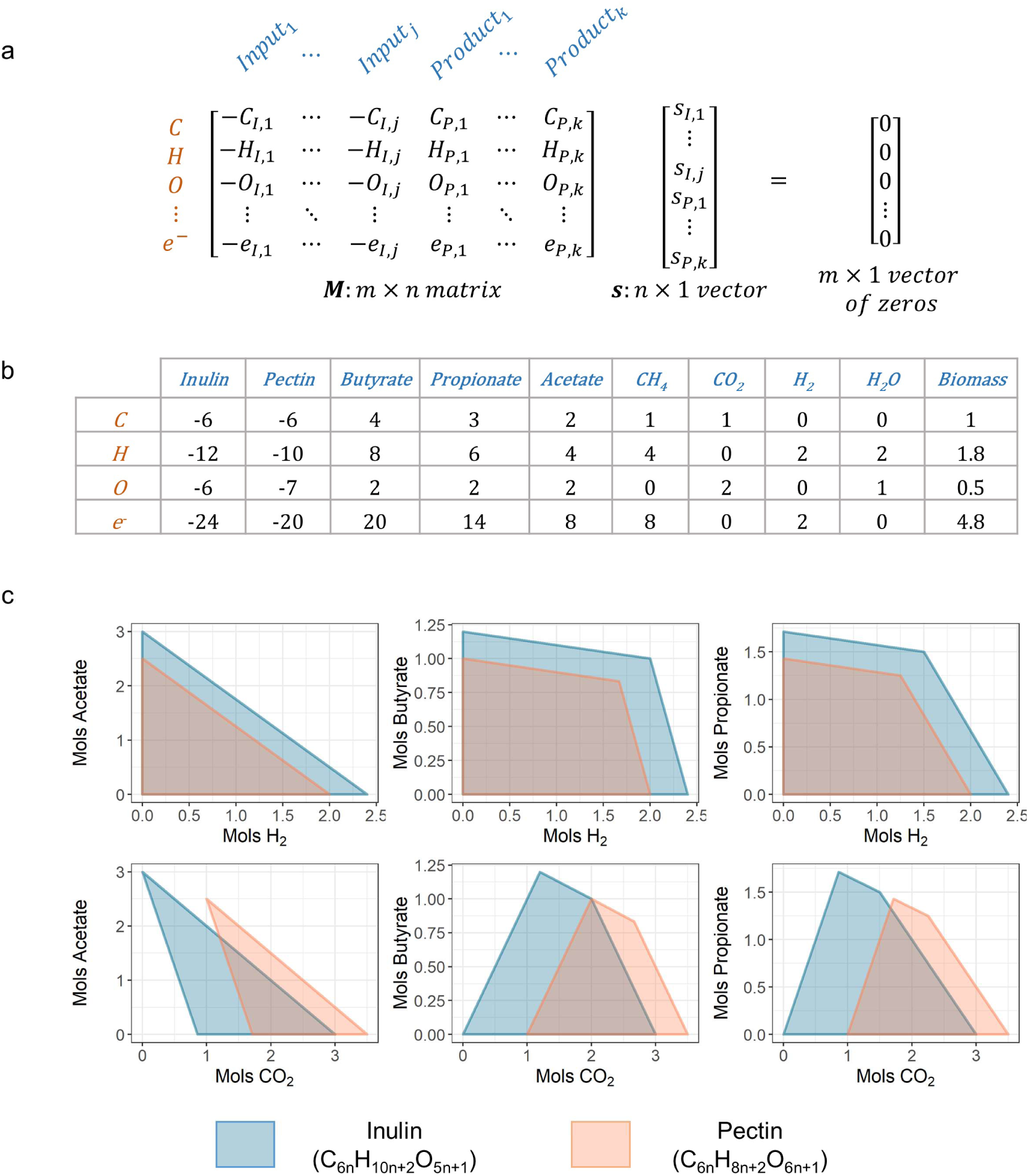
Modeling community production with mass and electron balance. a) A detailed representation of the theoretical model **Ms=0**, where **M** is a matrix with **n** chemicals (j inputs and k products). In **M**, inputs are represented in negative numbers while outputs are represented in positive numbers. Each row in **M** represents an element (or electrons) that needs to be balanced. b) The specific **M** matrix corresponding to our system of interest, fermentation of two different fibers, inulin and pectin. c) A set of selected 2D projections of the feasible product space predicted from our theoretical model for the fermentation of 1 mol inulin or 1 mol pectin. See Figure S1 for the full set of 2D projections for all fermentation products.

### Feasible product space of pectin fermentation is more limited compared to inulin

We applied our model to compare the feasible product space for the fermentation of 1 mol of inulin (C_6n_H_10n+2_O_5n+1_) to that of 1 mol of pectin (C_6n_H_8n+2_O_6n+1_) in a closed system. Since we were modeling product output from carbohydrate input, we only included C, H, O and valence electrons as rows in our matrix **M**. For products (columns) in **M**, we included the three most abundant SCFAs in the gut (acetate, propionate and butyrate), the three major components of intestinal gas (H_2_, CH_4_, and CO_2_), as well as water and biomass [represented by CH_1.8_O_0.5_N_0.2_, the mean chemical formula for microbial biomass (20), Figure 1b]. All product concentrations were restricted to be non-negative to simulate a closed system (*i.e.* product formation is solely from fiber input). Our model showed that in a closed system, the product space of pectin was more restricted than inulin (Figure S1); in particular, inulin had more potential for H_2_ production, while pectin had more potential for the production of CO_2_ (Figure 1c). The results of our model are in accordance with the simple intuition that a more oxidized substrate (pectin) would lead to more production of oxidized products such as CO_2_ and less production of reduced products such as H_2_. Model predictions were conserved even if further constraints were placed on the system, *i.e.* 15% of C in the fiber is converted into biomass as in a typical carbohydrate fermentation [Figure S2a, S2b, S2c (21)].

### Pectin degradation takes up reducing agents from the environment

We next asked if our theoretical predictions could be experimentally validated using an *ex vivo* framework in which we measured the response of stool microbiota to fiber addition. First, stool from 9 healthy human subjects was homogenized with phosphate saline buffer (PBS) under anaerobic conditions to create a fecal slurry. The slurry was then incubated in serum bottles at 37°C starting with 100% N_2_ in the headspace, with inulin, pectin, cellulose, or no additional fiber input (Figure 2a). Since our preliminary testing showed that fiber degradation in this system was almost entirely complete within 24h (19), we used the gas and SCFA concentrations at 24h as the experimental product concentrations for comparison to those predicted from our theoretical models. Because the fecal slurry itself contained a certain amount of residue material from food digestion in the human body, even samples that did not receive additional fiber produced gas and SCFAs; the fermentation products of a certain fiber in a sample was thus determined as the difference between product concentrations measured in a sample which received additional fiber and which did not.

**Figure 2.**
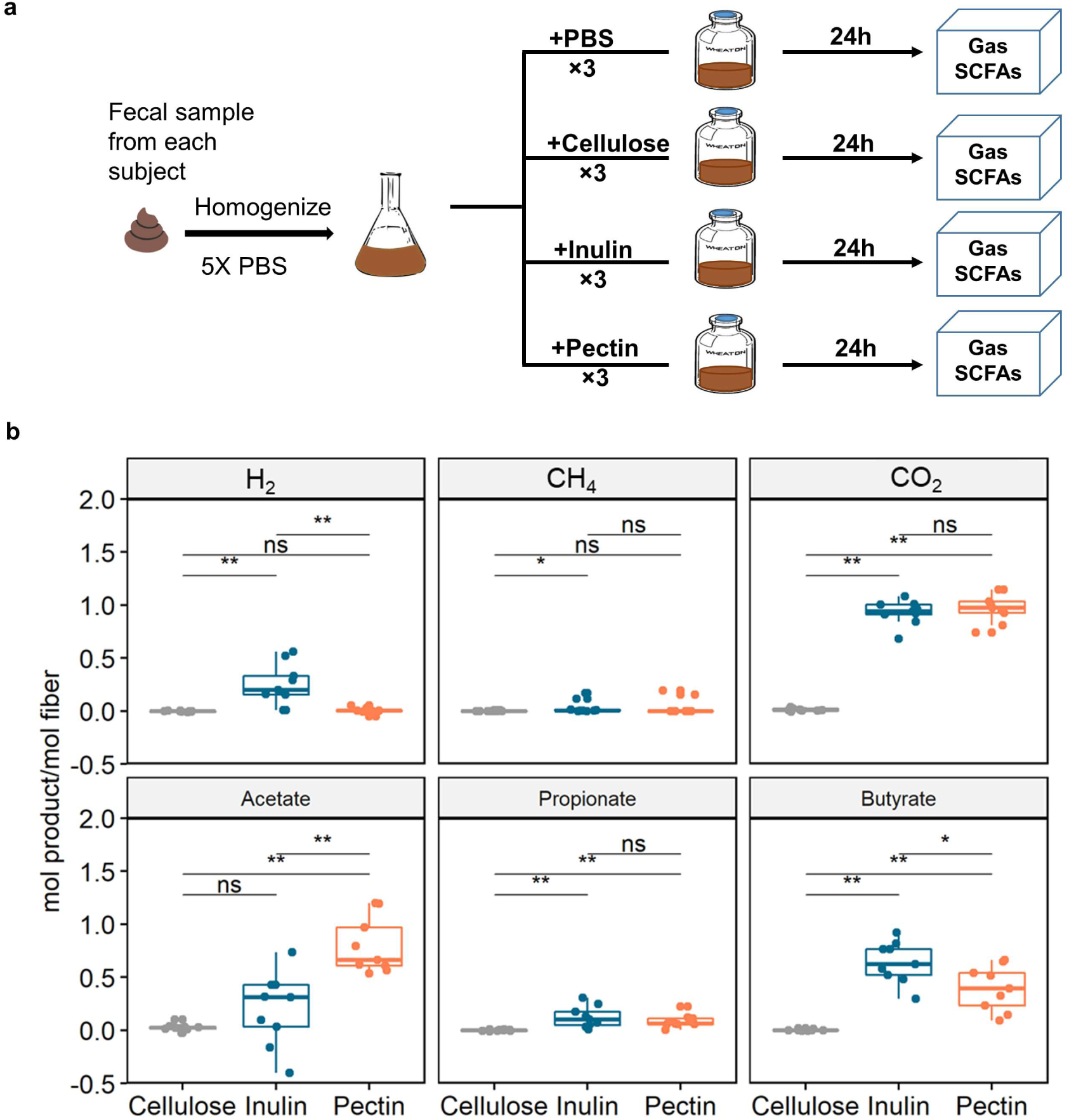
Inulin fermentation produces more H_2_ and less acetate than pectin in *ex vivo* system. a) Experimental scheme for studying fiber fermentation products in *ex vivo* system b) Major product concentrations in the *ex vivo* system after 24h measured as mol product production per mol of fiber. **, p<0.01, *, p<0.05 for paired Kruskal-wallis test.

Focusing on gas production in the *ex vivo* systems, we found that the amount of H_2_ produced by pectin fermentation was significantly lower than that of inulin (Figure 2b, Kruskal–Wallis test, paired, p=0.004), and the total amount of gas production was also lower (Kruskal–Wallis test, paired, p=0.07). However, we did not observe a higher amount of CO_2_ production in pectin fermentation compared to inulin fermentation as theoretically predicted; in fact, the measured CO_2_ productions from pectin fermentation did not fall within the previous theoretically predicted range (Figure 3a). This was also the case for acetate production in some samples that fermented inulin. Since our model only considers the most basic laws of chemistry and represents the maximum possible theoretical product range for a closed system, we hypothesized that the experimental violation of the results from the theoretical model was due to assuming that our experimental system was closed. Indeed, despite the serum bottle being a closed system with no material exchange with the environment outside of the bottle, fermentation of the additional fiber should be seen as a subsystem that can exchange products with the other subsystem in the bottle that ferments residue material in the fecal slurry.

**Figure 3.**
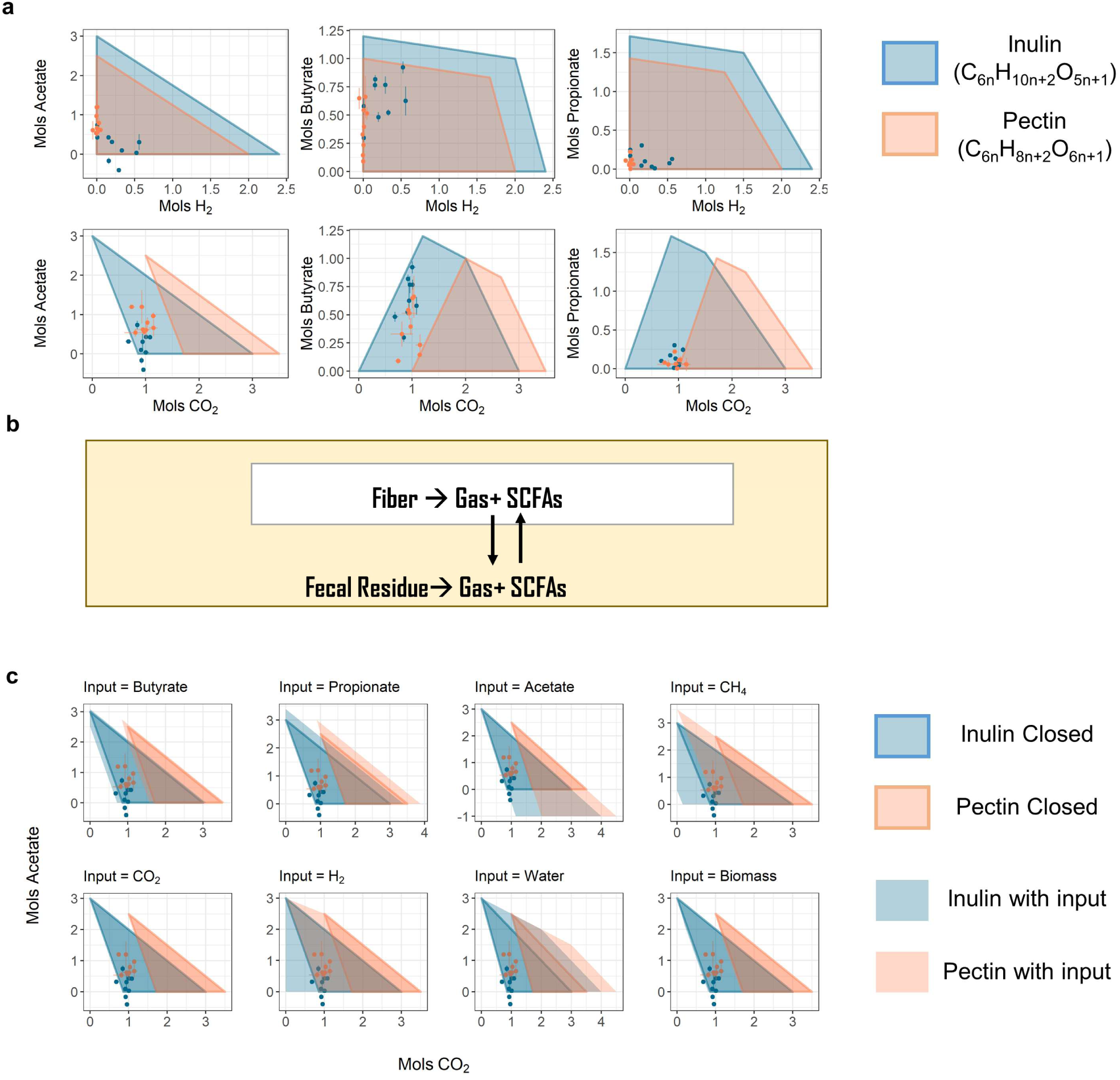
Pectin degradation requires the uptake of reducing agents. a) Comparison of product measurements in the *ex vivo* system to theoretically predicted feasible product ranges. b) Illustration of the two subsystems within the serum bottle and their material exchanges. c) Comparison of product measurements in the *ex vivo* system to theoretically predicted feasible product ranges in a closed system and when allowing different inputs.

We thus investigated what input the “fiber subsystem” would need from the “residue subsystem” for the measured CO_2_ to fall within the feasible product range determined by the theoretical model. Since on average the samples that did not receive additional fiber produced approximately 1/3 as much gas and SCFAs compared to those that did (Figure S4), we limited the input from the residue subsystem to the equivalent amount of product that can be produced by 1/3 mol of inulin or pectin. Allowing one input at a time, we found that only when H_2_ or CH_4_ was used as input would the measured CO_2_ fall within the predicted range (Figure 2c). Although we did not observe net uptake of either H_2_ or CH_4_ in our experimental data, but because both H_2_ and CH_4_ are chemicals with reducing power, there was likely influx of other reducing substrates not presented in our model from the “residue subsystem” to the “fiber subsystem”. Thus, in our *ex vivo* system, pectin degradation not only had a lower net production of H_2_ compared to inulin, but also took up reducing agents from the surrounding environment. We thus speculate that when pectin is degraded in the human gut, it is also taking up reducing agents—a process for which consequences are unclear and possibly worth further investigation.

### H_2_ and Acetate distinguishes the product profile of inulin and pectin degradation, but inter-personal variation of gas production is large

We next asked if the overall product profiles of inulin degradation and pectin degradation can be distinguished from each other, and whether changing fiber or microbial community contributed more to the variation in product profiles. We found that product profiles primarily clustered by fiber and not human subjects (Figure 4). Given that inulin fermentation generated significantly more H_2_ and less acetate compared to pectin (Figure 2b), we hypothesized these are the two major products that allowed the product profiles of inulin and pectin fermentation to be distinguished. Indeed, after training a Random Forest Classifier to predict whether a product profile was the result of inulin or pectin fermentation (AUC=1, p=0.03, paired t-test, two-sided), we found that H_2_ and acetate were the two most important features for predicting the fiber of fermentation (Figure S3). Not only did the oxidation state of the fiber have impact on gas production, it also influenced SCFA production: the more oxidized substrate, pectin, produced more of the most oxidized SCFA, acetate. Also, aside from H_2_, inulin also produced more of the most reduced SCFA, butyrate (Figure 2b).

**Figure 4.**
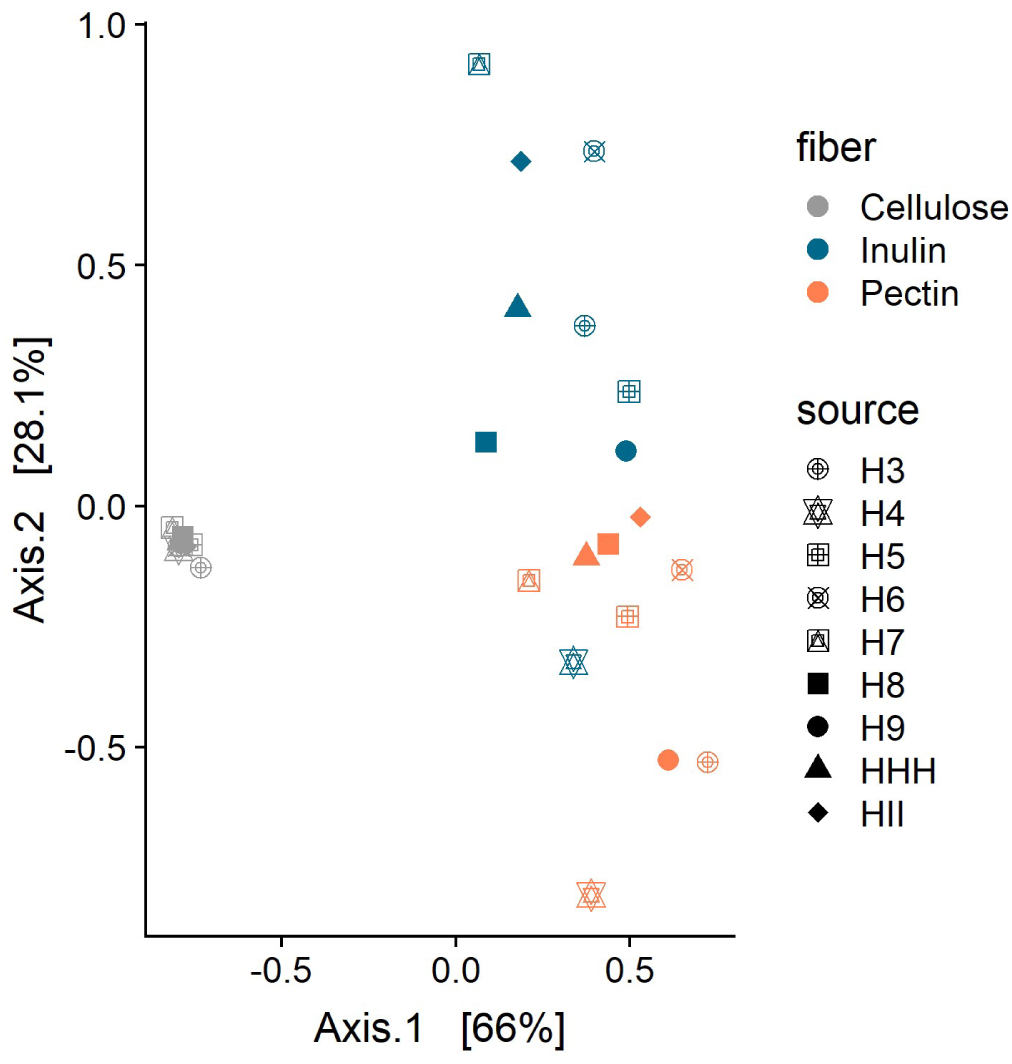
Principle coordinate analysis (PCoA) of the product profiles of different fibers.

Having determined that fiber type strongly impacts product profiles, we next investigated inter-individual differences. Euclidian distances between the product profiles of different fibers within the same person were not significantly larger than those between different people on inulin (Mann–Whitney U test, p=0.25), and slightly larger for pectin (Mann–Whitney U test, p=0.03). Thus, the microbiota of different people exhibits functional heterogeneity in converting the same fibers into products of different quantities and composition.

### Relative effects of microbiome and substrate chemistry on gas production differ among gases

We further explored if there were signatures within the microbiomes that promoted the production of gases. Since levels of H_2_ production were generally low for pectin fermentation, we investigated if there were specific ASVs associated with net H_2_ production during inulin fermentation. Selecting for these ASVs via Lasso regression identified a *Lachnospiraceae* amplicon sequencing variant (ASV) positively associated with net H_2_ production (Figure 5a, Pearson’s r=0.95, p= 6.2×10^−5^). Since net H_2_ production in the gut is the difference between the total production of H_2_ and the total consumpution of H_2_ (12), and *Lachnospiraceae* can be either hydrogen producers or consumers (22, 23), the positive association of the *Lachnospiraceae* ASV with H_2_ production in the human gut may indicate that net H_2_ production is more dependent on H_2_ production than consumption. Consistent with this hypothesis, H_2_ consumption abilities of gut microbiota may be more consistent between different people compared to H_2_ production because of the higher diversity of H_2_ consumption pathways (methanogenesis, reductive acetogenesis and sulfate reduction) compared to production. It is, however observed that net H_2_ production was lowest in the samples that produced methane, probably because the amount of sulfate in the *ex vivo* system is not enough to for the most energetically favorable H_2_ consumption pathway, sulfate reduction, to consume all the H_2_ produced (Supplementary text), and methanogenesis is more energetically favorable than reductive acetogenesis.

**Figure 5.**
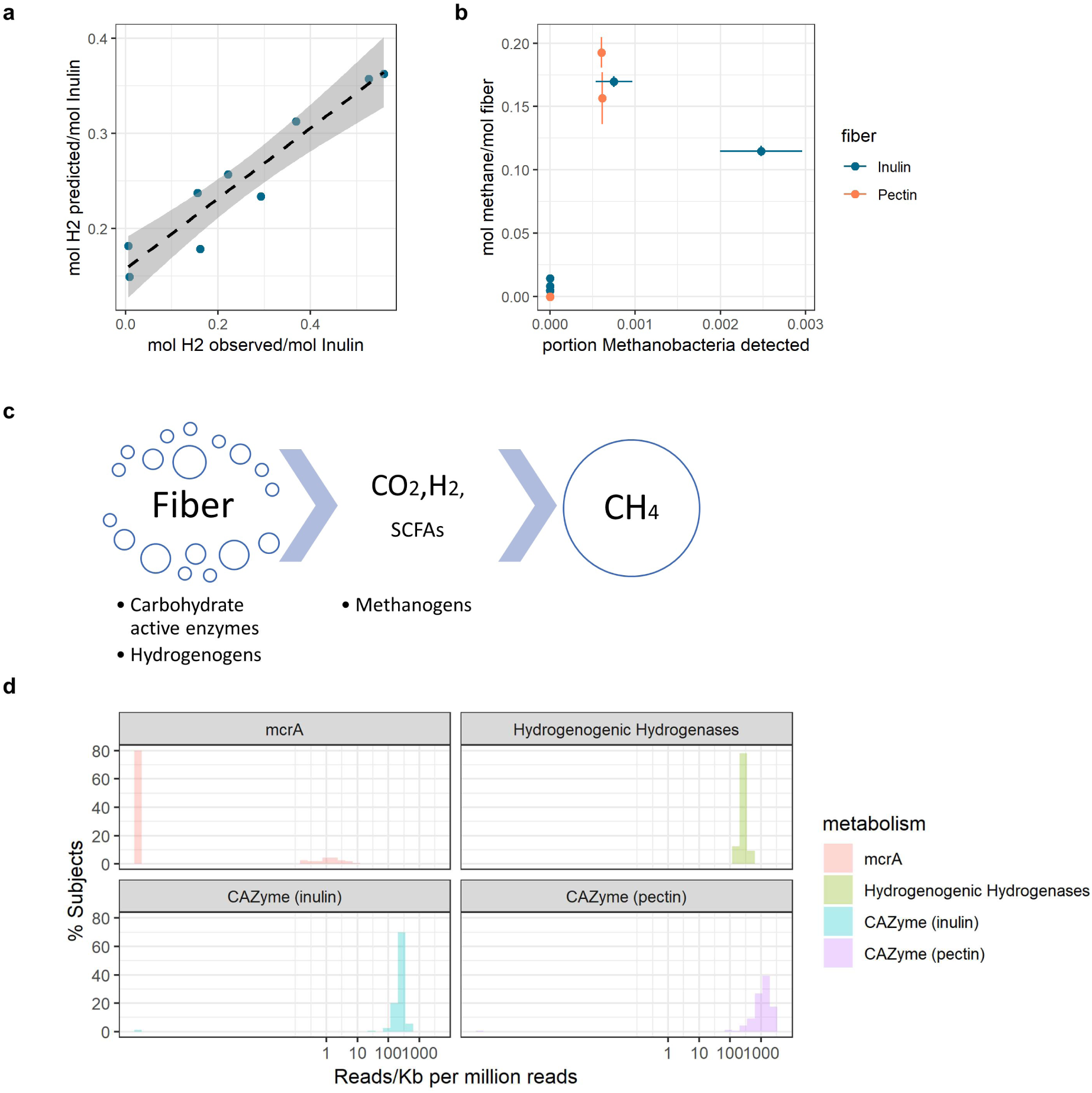
Different gases are differentially influenced by substrate chemistry and gut microbiome composition. a) Relationship between the predicted value of H_2_ production per mol of inulin based on a linear model with a *Lachnospiraceae* ASV as the independent variable, and the observed amount of H_2_ production per mol of inulin in *ex vivo* system. Dotted line represents a linear fit of the relationship, with grey areas around the line representing the standard error of the fit. b) Relationship between the production of methane per mol of fiber in the *ex vivo* system and the relative abundance of *Methanobacteria* in the samples. Error bars represent standard deviation (n=3). c) Schematic of fiber degradation and production of gas and SCFAs. d) Distribution of the abundance of the methanogenesis marker gene mcrA, hydrogenenic hydrogenases, and CAZymes in the metagenomes of 160 different people in the HMP dataset. All gene counts were increased by 10^−5^ so that the log-scaled x-axis could accommodate samples with zero hits.

In contrast to H_2_, whose production was strongly affected by substrate, we found that methane production was solely dependent on whether there were detectable levels of *Methanobacteria* in the microbiota (Figure 5b). Given that methane is a downstream product of H_2_ (Figure 5c), we asked why methane production was not affected by substrate chemistry as H_2_ production was. We hypothesized that this is because there are generally large amounts of bacteria in the human gut that contain carbohydrate active enzymes and hydrogenases that allow fiber breakdown and hydrogen production; thus the amount of these enzymes are not a limiting factor, allowing hydrogen production to be instead dependent on substrate stoichiometry. Meanwhile, a large portion of the human population do not harbor sufficient numbers of methanogens, making them the limiting factor for methane production; however, when the number of methanogens are sufficient they are not limited by the amount of H_2_ production because methanogens are stronger competitors for H_2_ compared to reductive acetogens. To test this hypothesis, we surveyed the abundance of carbohydrate active enzymes (CAZymes) for inulin and pectin, as well as hydrogenogenic hydrogenases, and methyl-CoM reductases (mcrA, marker gene for methanogens) in the metagenomes of 160 randomly selected healthy human subjects from the human microbiome project (HMP). While only approximately 20% of subjects had detectable levels of methanogens, nearly all subjects harbored CAZymes for inulin and pectin degradation as well as hydrogenogenic hydrogenases (Figure 5d). The percentage of subjects (20%) with detectable methanogens in the HMP data is in accordance with our results: 2 out of the 9 subjects were methane producers in our *ex vivo* experiment. Thus, overall, the production of more “general” metabolites such as H_2_ is more likely to be affected by the chemical composition of the prebiotic, while more “rare” metabolites such as methane are more likely to be limited by the organisms that produce it.

## Discussion

In this study, we used a combination of theoretical models and an *ex vivo* experimental framework to examine how the chemistry of prebiotics and the composition of the gut microbiota influence gas production during prebiotic fermentation by gut microbiota. Specifically selecting two different common prebiotics (inulin and pectin) with different levels of oxidation, we find that metabolites that can be produced by more organisms in the human gut, such as H_2_, are more affected by the chemical composition of prebiotics compared to metabolites that are produced by less common organisms in the gut, such as methane. Overall, these results suggest that both the chemical nature of the prebiotic and the individual’s gut microbiome needs to be taken into account when administering prebiotics to individuals.

Our data also reveal that there may be general trade-offs in the production of SCFAs verses gas. For example, while inulin fermentation leads to more production of the more reduced SCFA butyrate, it also leads to more production of the reducing agent H_2_, and in turn increasing overall gas production. However, which is more preferable for the subject—more production of butyrate, or less production of overall gas, or just less production of H_2_ is often unknown and specific to the individual subject. This can be further complicated if inter-individual differences in H_2_ and SCFA production is considered: not every individual produces more butyrate when fermenting inulin. Similarly, for pectin we were able to infer by comparing the experimental data to the theoretical product range that pectin degradation requires uptake of reducing agents from the surrounding environment. Again, what effect this has on the host is unknown: would the uptake of these reducing agents lead to the generation of more reactive oxygen species that can directly attack cells in the gut epithelial barrier, interfere with iron uptake, or initiate lipid peroxidation processes (24)? Would this be costlier to the host compared to generating more H_2_? More importantly, we also lack a way to evaluate if the scale of the differences is large enough for them to count as a factor in prebiotic selection.

These problems emphasize that the effect of prebiotics on gut and human health must be looked at from both individual and systems perspectives. Often a compound is deemed as a prebiotic because it can increase the growth of known beneficial microbes such as Bifidobacteria and Lactobacilli, or promote the production of target metabolites. However, the full diversity of a mixed culture environment such as the human gut must be considered when selecting for prebiotics: it is very hard to only selectively grow organisms that produce one or a few metabolites of interest, and the effect of any by-products must be considered. Inter-individual differences in product formation due to heterogeneity in gut microbiota composition, as well as responses to the metabolites produced, must also be considered. Our use of theoretical modeling and the *ex vivo* experimental system to explore gas production and its relationship to SCFA production is just a beginning: these are relatively cheap and simple methods to shine light on important points that should be considered in prebiotics design. In the future, a more systematic evaluation on what important factors other than the formation of beneficial metabolites should be considered in prebiotic design is needed.

## Methods

### Experimental model and participant details

9 healthy human volunteers were enrolled into the study under the supervision of the MIT Committee on the Use of Humans as Experimental Subjects (COUHES), who approved the study under protocol number 1510271631. All participants provided written, informed consent, and the study was conducted in accordance with the relevant guidelines and regulations. To be included, participants had to be between 18-70 years of age, have a BMI between 18 and 30, and do not have a history of inflammatory bowel diseases/syndrome, Type-2 diabetes, kidney diseases, intestinal obstruction, or colorectal cancer. They were also not currently pregnant, breast-feeding or have received antibiotics treatment in the 6 months leading up to the study. Enrollment occurred between June 2017 and Oct 2017. The study group included 4 females and 5 males, all between 25-40 years of age.

### Linear systems model for modeling community production

The product space for the system **Ms=0** is a polytope defined by linear combinations of the basis vectors of Null(M), i.e. **Bx=s**. Constraints on the product space (i.e. for the closed system all elements in s corresponding to products are non-negative) were used to find the vertices of the polytope of **x** by converting the half-space representation (the intersection of half spaces, represented by **Bx=s**) into vertex representation (set of extreme points of the polytope). Vertices of the polytope of **s** were calculated from multiplying **B** with the vertices of polytope **x**. The vertices for **s** were used to draw 2D hulls for pairs of products to visualize the product polytope, as in Figure 1c, 3a and 3c. When product input was allowed for the system, the constraint on the element in **s** corresponding to the input product would be relaxed to be larger than the negative of the equivalent amount of product that can be produced by 1/3 mol of inulin or pectin (on average control samples produced approx. 1/3 as much SCFA and/or gas than samples with inulin or pectin treatment, Figure S4).

The basis set of vectors for the null space of matrix M was calculated from the QR-decomposition of the matrix using the R package “pracma”(25). The conversion of half-space representation to vertex representation of polytopes were performed using the R package “rcdd”(26).

### Setup of *ex vivo* system

The setup of the *ex vivo* system was the same as in Gurry et al 2020 (19), with some adaptation for gas measurements. Briefly, fresh stool samples were collected and homogenized with reduced PBS containing 0.1% L-Cysteine in a ratio of 1g/5ml. Fiber was spiked in to the homogenates from stock solutions such that the final concentrations of fibers in the samples were as follows: control no fiber, 10g/L inulin, 5g/L pectin or 20g/L cellulose. For each participant, 2mL of the final fecal slurry of each condition was added in triplicates to 60mL glass serum bottles (Supelco, Bellefonte PA). The serum bottles containing the samples were transferred to a vinyl anaerobic chamber filled with 100% N_2_, with no detectable amounts of CO_2_ and H_2_, and sealed in the chamber using magnetic crimp seals with PTFE/silicone septa (Supelco, Bellefonte PA). A total of 12 bottles per participant were incubated at 37°C for 24h with no shaking.

### Gas and SCFA measurements

Concentrations of headspace gases were determined using gas chromatography. We used a Shimadzu GC-2014 gas chromatography (GC) configured with a packed column (Carboxen-1000, 5’ x 1/8” (Supelco, Bellefonte PA)) held at 140°C, argon carrier gas, thermal conductivity (TCD) and methanizer-flame ionization detectors (FID). At the end of the 24h incubation period, subsamples of the headspace (0.20 cm^3^ at the laboratory temperature, *ca*. 23°C) from each serum bottle were taken via a gas-tight syringe and injected onto the column. Gas concentrations were determined by comparing the partial pressures of samples and standards with known concentrations. Accuracy of the analyses, evaluated from standards, was ±5%. Measurements of H_2_ were taken using the TCD while measurements of CH_4_ and CO_2_ were taken with the FID.

SCFA measurements were made from taking 1mL of fecal slurry from each serum bottle immediately after the GC measurements were taken, and freezing the fecal slurry at -80°C until time of measurement. SCFA measurements were made on an Agilent 7890B system with a FID at the Harvard Digestive Disease Core (Agilent Technologies, Santa Clara, CA). Detailed procedures SCFA measurements are the same as in Gurry et al 2020 (19). Although the amount of 10 volatile acids (acetic, propionic, isobutyric, butyric, isovaleric, valeric, isocaproic, caproic, and heptanoic acids) were reported, all but the acetic, butyric and propionic acids were in trace amounts and we only used these three SCFAs for our models.

### Machine learning and Statistics

The Principle Coordinate Analyses of the *ex vivo* fermentation products were performed with the R package ‘ape’ using Euclidian distance matrices (27). The Random Forest Classifier of product profiles were built using the R package “randomForest” [(28), ntree=2000)]. The samples were apportioned into test and training sets with a 50-50 split generated by a random seed. The ROC and AUC of the classifier were calculated using the R package “pROC”(29), and the p-value of the AUC was determined by comparing the classifier AUC to the AUC calculated when fiber categories of the test set sampled were randomly assigned. Lasso regression was performed with the R package “glmnet” (30) and cross-validation was performed with a leave-one-out approach.

### DNA extraction, library prep and sequencing

The MoBio PowerSoil-htp 96 kit (now Qiagen Cat No./Id: 12955-4), with minor modifications, was used to extract the DNA from all fecal samples. For all samples, 250uL of the fecal slurry was used with the MoBio High Throughput PowerSoil bead plate (12955-4 BP). 16S rRNA gene amplicon libraries (V4 hypervariable region, U515-E786) using a two-step PCR approach were prepared according to the method described in Preheim *et al* (31). Samples were sequenced on an Illumina MiSeq (PE 150+150) at the Broad Walk-up Sequencing platform (Broad Institute, Cambridge, MA). The average sequencing depth of the samples were 53,046 reads/sample.

### 16S rRNA amplicon data analysis

All 16S rRNA amplicon libraries were processed according to a custom pipeline based on DADA2, as described in Yu *et al* (32), except that only the forward reads were used due to issues in merging reads. The output of the pipeline was amplicon sequencing variants (ASVs). Taxonomy for all sequence variants was assigned using the RDP database.

### Metagenome analysis

We downloaded 160 randomly selected metagenomes from the human stool microbial communities of the Human Microbiome Project (National Institutes of Health, USA). Each metagenome was rarefactioned to 20 million reads (forward+reverse) using seqtk seeded with the parameter –s100. The rarefactioned metagenomes were screened in DIAMOND (maximum number of high scoring pairs (HSPs) per subject sequence to save for each query=1, blastx) against hydrogenogenic hydrogenases retrieved from the HydDB database (33), mcrA genes retrieved from the PhyMet database (34), and CAZymes from the dbCAN database (35). Results were then filtered [length of amino acid>25 residues, percent identical matches>65% (mcrA and hydrogenases) or >35% (CAZymes)]. Reads were eventually normalized to reads per kilobase million using the formula 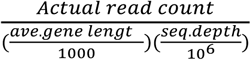.

## Acknowledgements

This work was supported by a grant from the Rasmussen Family Foundation. We are especially highly thankful to the Ono Lab at MIT and Jeemin Rhim for teaching and allowing us to use their GC for measurements of gas production, and David Sontag at the MIT Computer Science and Artificial Intelligence Laboratory for consultation with the linear systems model. We thank the Broad walk-in sequencing center for their assistance with sequencing, and Harvard Digestive Disease Core for their assistance on GC for measurements of SCFA production. We also wish to thank Chengzhen Dai, Siavash Isazadeh, Chuliang Song, Fangqiong Ling, and Martin Polz for helpful discussions and/or comments on various versions of this manuscript.

## Data availability

All amplicon sequencing data generated in this study can be accessed upon publication on the US National Center for Biotechnology Information SRA database under BioProject PRJNA587309. All gas and SCFA measurements, ASV tables, and code for data analysis will be available at https://github.com/cusoiv.

---

**Figure S1.**
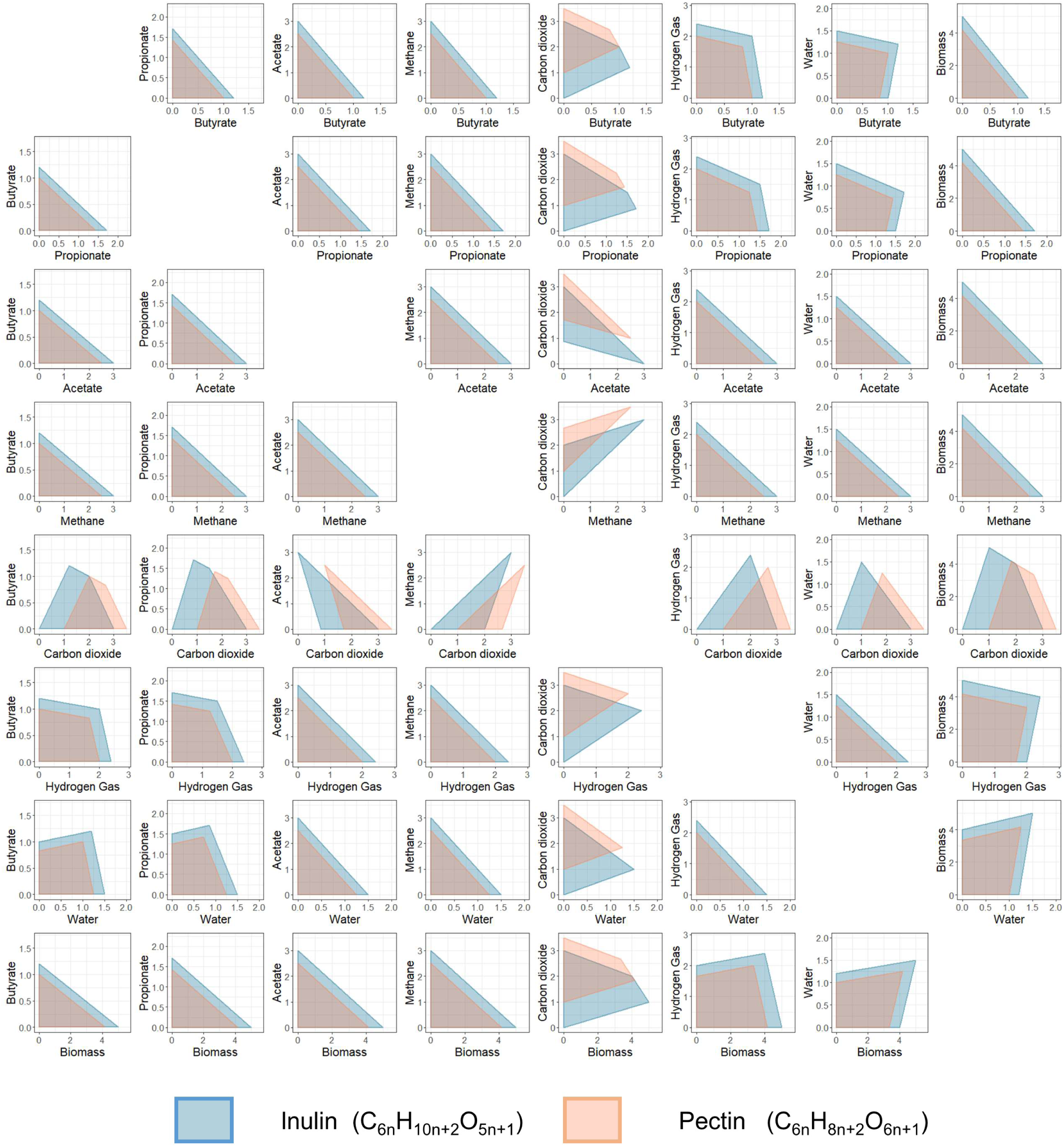
2D projections of the feasible product space for all fermentation products. The full set of 2D projections of the feasible product space predicted from our theoretical model for the fermentation of 1 mol inulin or 1 mol pectin. Units on all axes are mol product/mol fiber.

**Figure S2.**
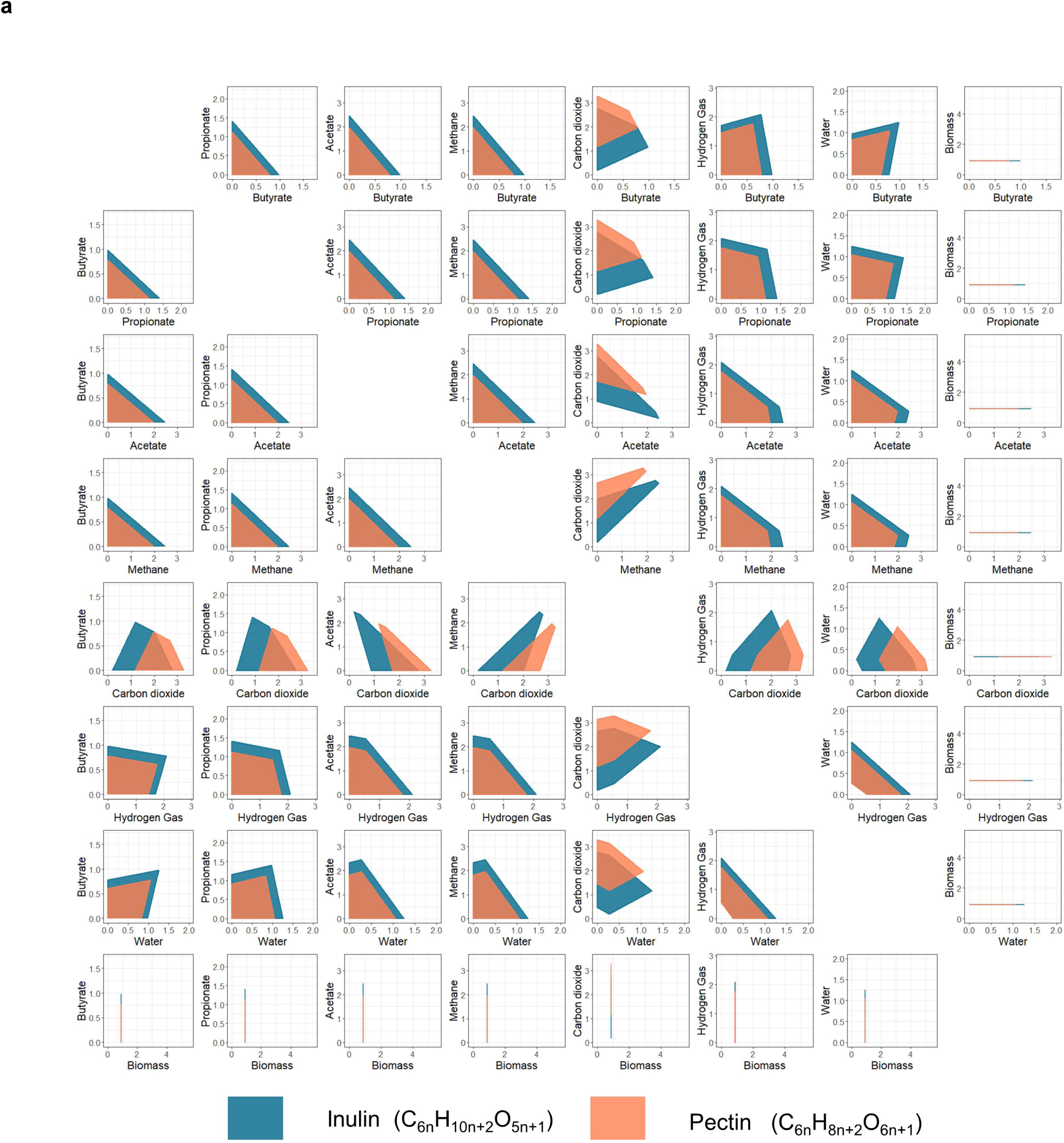
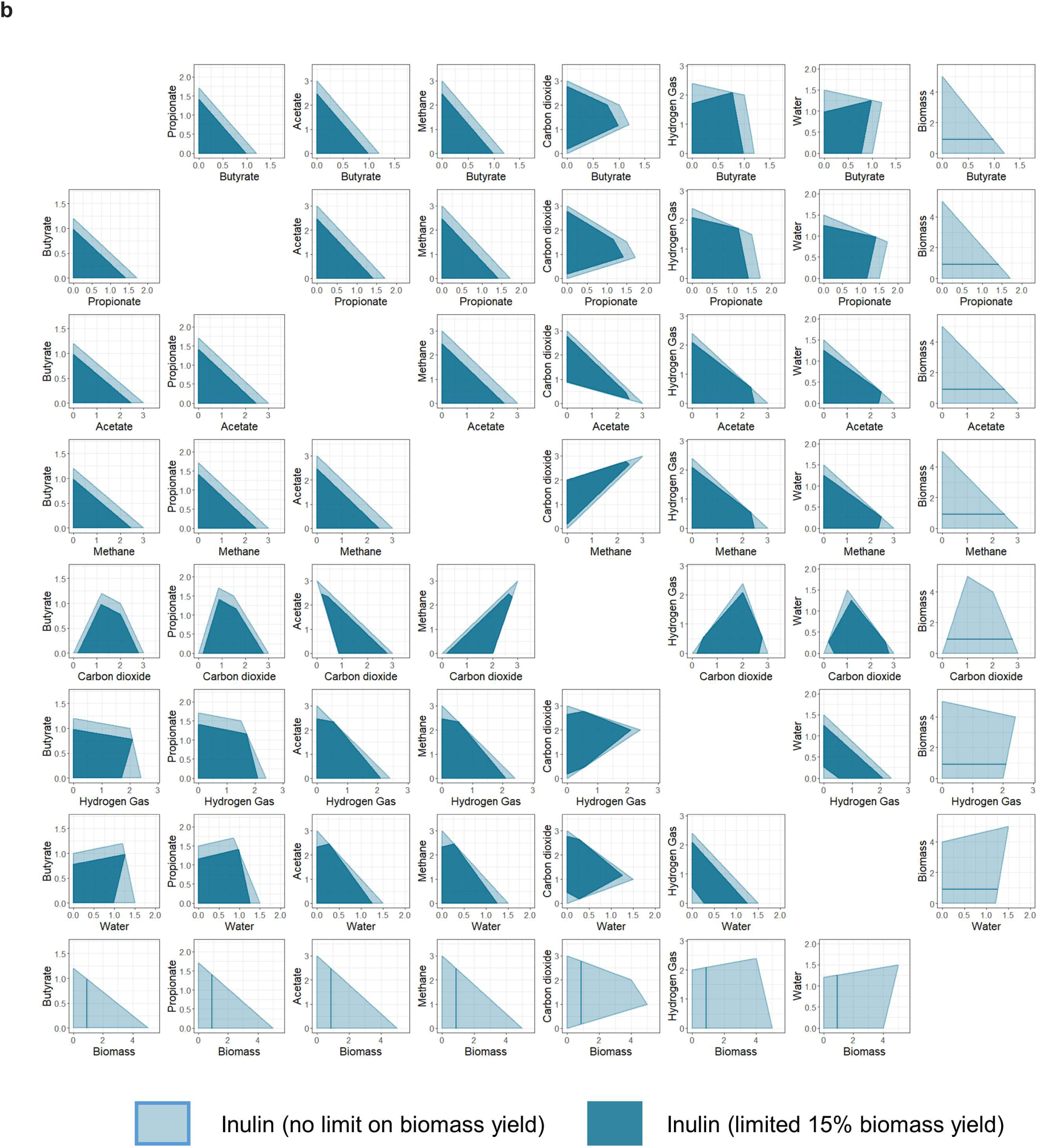
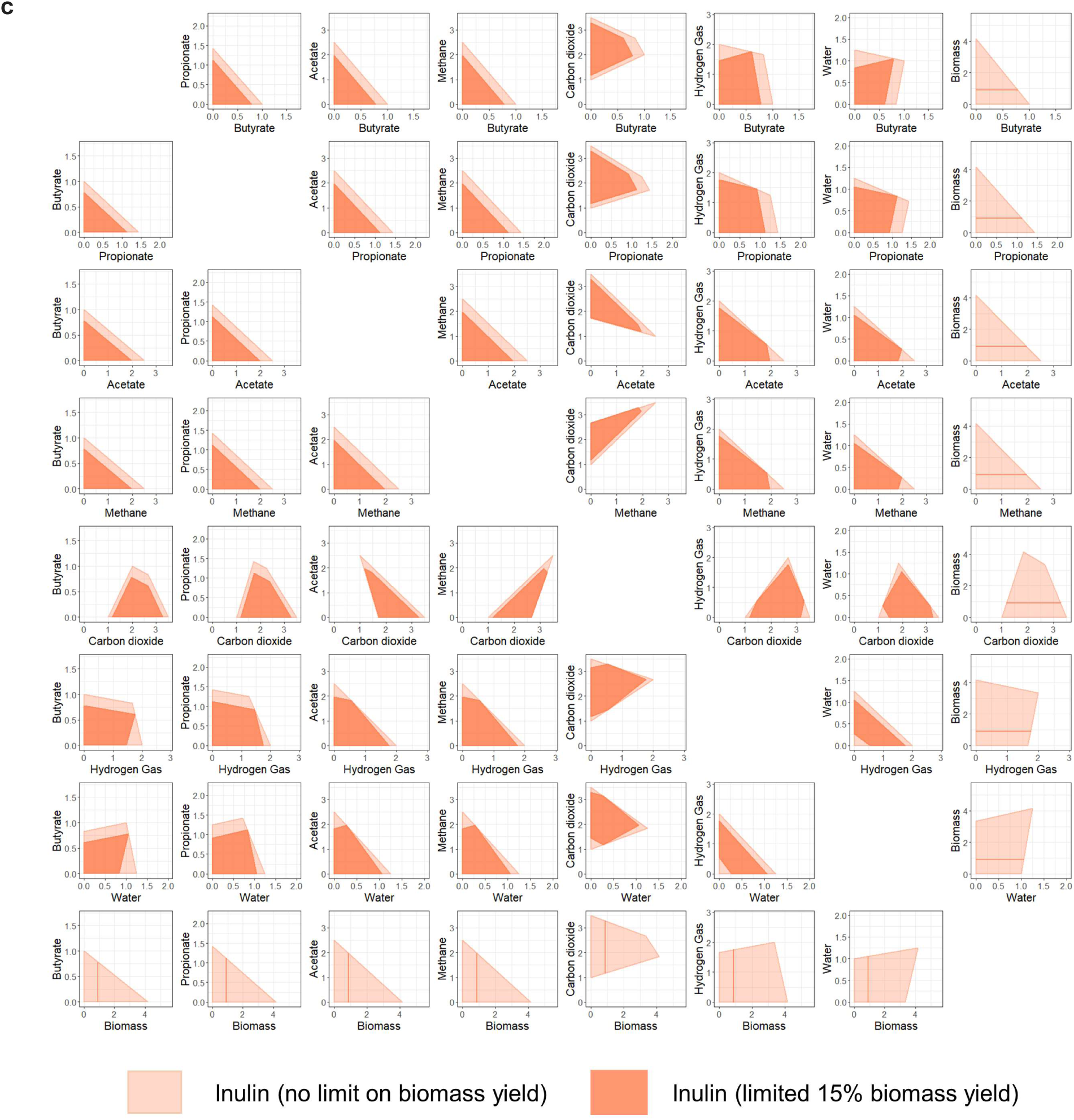
2D projections of the feasible product space for all fermentation products assuming constant biomass yield. a) The full set of 2D projections for the feasible product space predicted from our theoretical model for the fermentation of 1 mol inulin or 1 mol pectin, when assuming that 15% of the C in fiber is eventually converted into biomass. Units on all axes are mol product/mol fiber. b) Comparison of the 2D projections for the feasible product space predicted from our theoretical model for 1 mol of inulin with no restriction on biomass yield (light blue) and assuming a 15% biomass yield during the fermentation process (dark blue). c) Comparison of the 2D projections for the feasible product space predicted from our theoretical model for 1 mol of pectin with no restriction on biomass yield (light coral) and assuming a 15% biomass yield during the fermentation process (coral).

**Figure S3.**
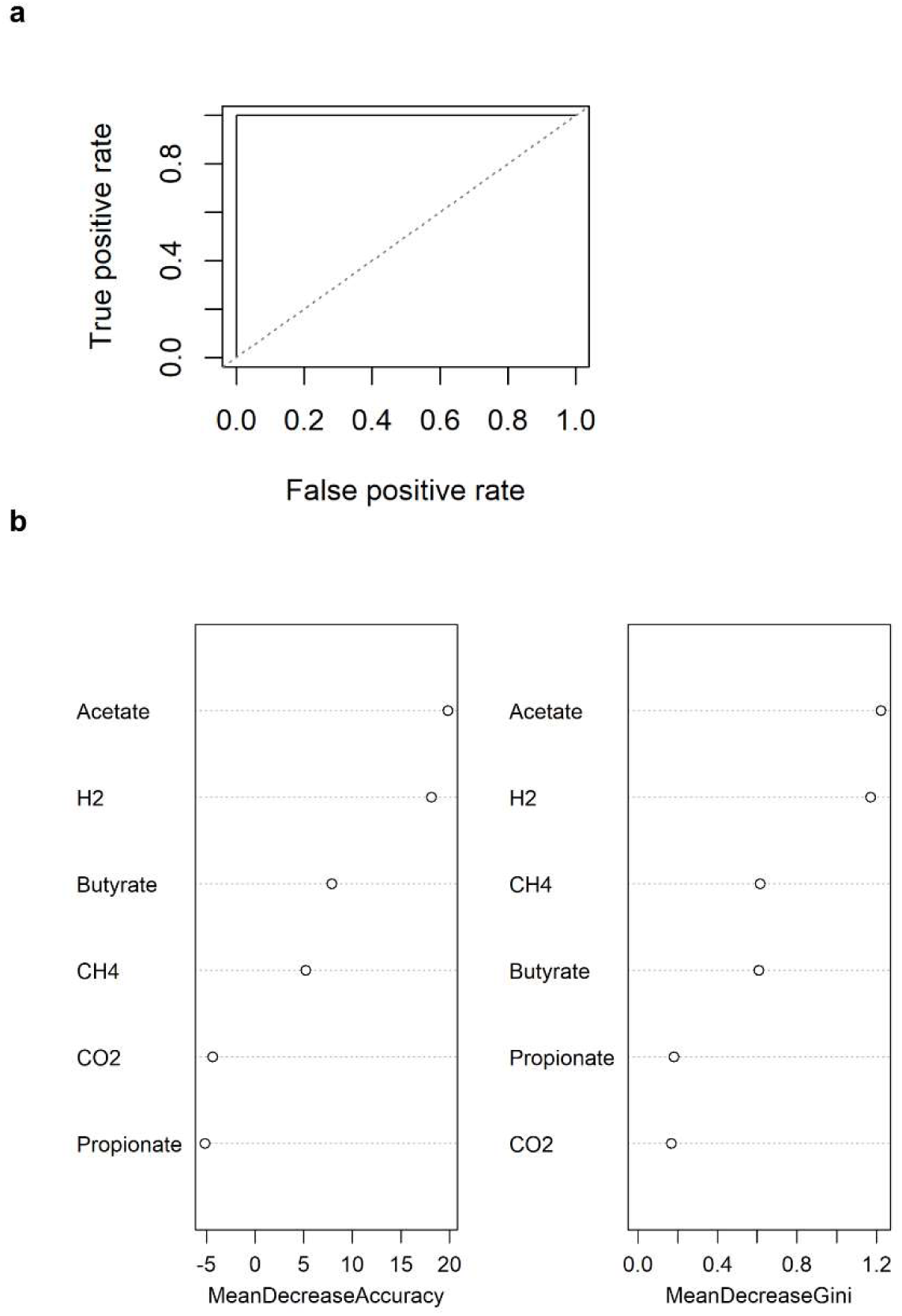
Performance of Random Forest Classifier for predicting prebiotic substrate based on product profile. a) Receiver operating characteristic (ROC) curve of the classifier b) Importance graphs for the classifier, ranked by top to bottom by two parameters (Mean Decrease Accuracy or Mean Decrease Gini) that represent the increase in misclassification produced on average by removing the given predictor (more important predictors are on top).

**Figure S4.**
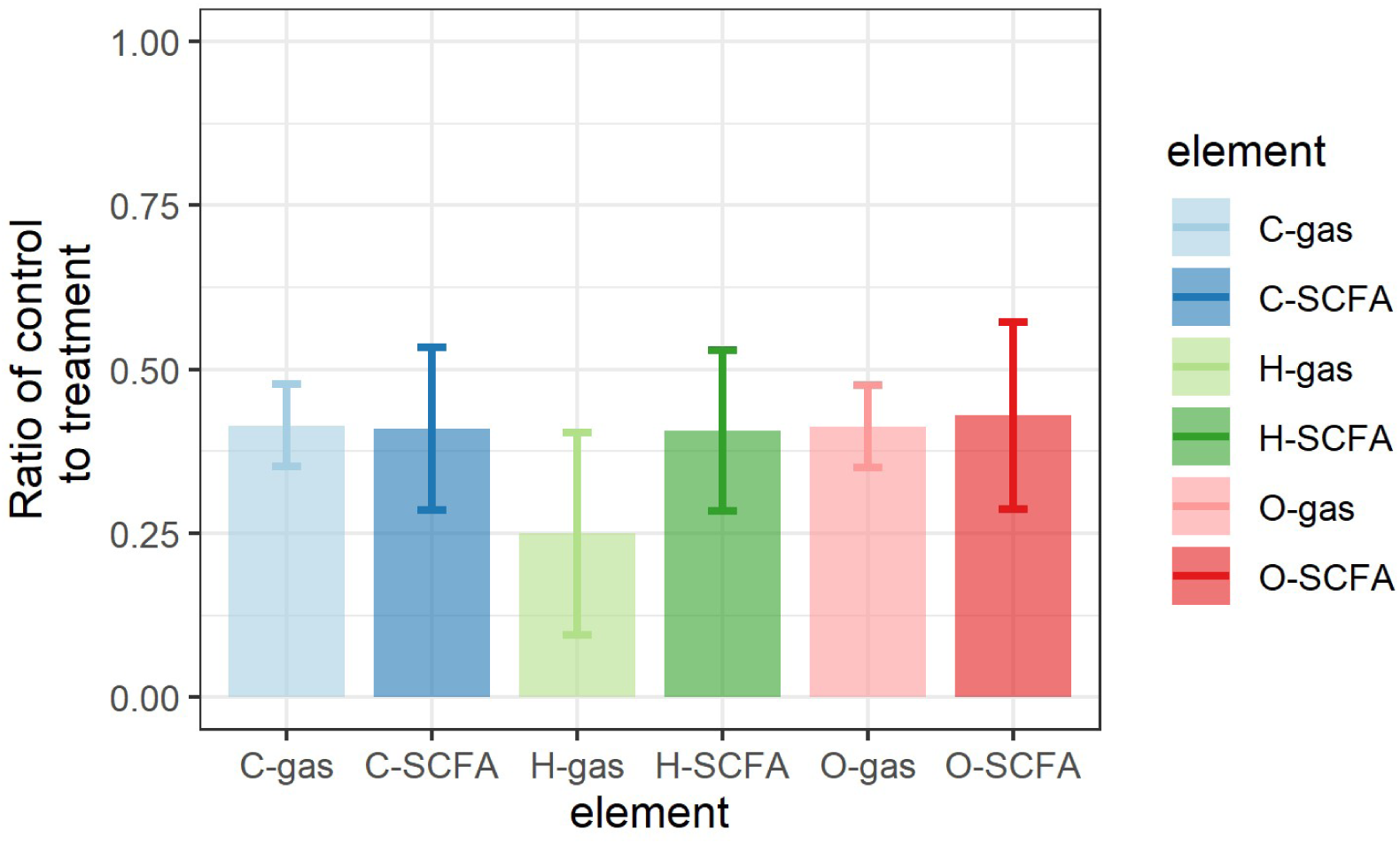
Ratio of products in control versus treatment groups. The gas or SCFAs in each sample were summed together by their Carbon (C), Hydrogen (H) or Oxygen (O) content. Only the samples with inulin or pectin addition were counted in the treatment group.

## Supplementary Text

### Estimation of Sulfate content in *ex vivo* system

The content of sulfate in human feces is estimated to be 2.7µmol/g, independent of diet (36). Since each serum bottle in our ex vivo system contains approximately 0.4g of feces, the sulfate content in each bottle is 1.08µmol, which would take 4.32umol of H_2_ to undergo complete reduction to hydrogen sulfide. This is lower/approximately the same compared to the net amount of H_2_ left in the serum bottles at our time of measurement (32±22 µmol for inulin, 3.2±1.4 µmol for pectin), and given that gross production should be not less than net production, it is likely that the amount of sulfate the *ex vivo* system does not support elimination of H_2_ based on sulfate reduction alone.

## References

1. Cotillard A, Kennedy SP, Kong LC, Prifti E, Pons N, Le Chatelier E, Almeida M, Quinquis B, Levenez F, Galleron N, Gougis S, Rizkalla S, Batto J-M, Renault P, ANR MicroObes Consortium, Doré J, Zucker J-D, Clément K, Ehrlich SD, ANR MicroObes consortium Members, Blottière H, Leclerc M, Juste C, de Wouters T, Lepage P, Fouqueray C, Basdevant A, Henegar C, Godard C, Fondacci M, Rohia A, Hajduch F, Weissenbach J, Pelletier E, Le Paslier D, Gauchi J-P, Gibrat J-F, Loux V, Carré W, Maguin E, van de Guchte M, Jamet A, Boumezbeur F, Layec S. 2013. Dietary intervention impact on gut microbial gene richness. Nature 500:585–588.

2. Tremaroli V, Bäckhed F. 2012. Functional interactions between the gut microbiota and host metabolism. Nature 489:242–249.

3. De Filippis F, Vitaglione P, Cuomo R, Berni Canani R, Ercolini D. 2018. Dietary Interventions to Modulate the Gut Microbiome—How Far Away Are We From Precision Medicine. Inflamm Bowel Dis 24:2142–2154.

4. Gibson GR, Hutkins R, Sanders ME, Prescott SL, Reimer RA, Salminen SJ, Scott K, Stanton C, Swanson KS, Cani PD, Verbeke K, Reid G. 2017. Expert consensus document: The International Scientific Association for Probiotics and Prebiotics (ISAPP) consensus statement on the definition and scope of prebiotics. Nat Rev Gastroenterol Hepatol 14:491–502.

5. Holscher HD. 2017. Dietary fiber and prebiotics and the gastrointestinal microbiota. Gut Microbes 8:172–184.

6. Canani RB, Costanzo MD, Leone L, Pedata M, Meli R, Calignano A. 2011. Potential beneficial effects of butyrate in intestinal and extraintestinal diseases. World J Gastroenterol WJG 17:1519–1528.

7. Belcheva A, Irrazabal T, Martin A. 2015. Gut microbial metabolism and colon cancer: Can manipulations of the microbiota be useful in the management of gastrointestinal health? BioEssays 37:403–412.

8. Shibata N, Kunisawa J, Kiyono H. 2017. Dietary and Microbial Metabolites in the Regulation of Host Immunity. Front Microbiol 8.

9. Míguez B, Gómez B, Gullón P, Alonso BG and JL. 2016. Pectic Oligosaccharides and Other Emerging Prebiotics. Probiotics Prebiotics Hum Nutr Health.

10. Verspreet J, Damen B, Broekaert WF, Verbeke K, Delcour JA, Courtin CM. 2016. A Critical Look at Prebiotics Within the Dietary Fiber Concept. Annu Rev Food Sci Technol 7:167–190.

11. Müller V. 2008. Bacterial FermentationeLS. American Cancer Society.

12. Carbonero F, Benefiel AC, Gaskins HR. 2012. Contributions of the microbial hydrogen economy to colonic homeostasis. Nat Rev Gastroenterol Hepatol 9:504–518.

13. Cormier RE. 1990. Abdominal Gas, p.. In Walker, HK, Hall, WD, Hurst, JW (eds.), Clinical Methods: The History, Physical, and Laboratory Examinations, 3rd ed. Butterworths, Boston.

14. Azpiroz F. 2005. Intestinal gas dynamics: mechanisms and clinical relevance. Gut 54:893–895.

15. Sahakian AB, Jee S-R, Pimentel M. 2010. Methane and the Gastrointestinal Tract. Dig Dis Sci 55:2135–2143.

16. Seo AY, Kim N, Oh DH. 2013. Abdominal Bloating: Pathophysiology and Treatment. J Neurogastroenterol Motil 19:433–453.

17. McIntosh K, Reed DE, Schneider T, Dang F, Keshteli AH, De Palma G, Madsen K, Bercik P, Vanner S. 2017. FODMAPs alter symptoms and the metabolome of patients with IBS: a randomised controlled trial. Gut 66:1241–1251.

18. Gibson PR, Shepherd SJ. 2010. Evidence-based dietary management of functional gastrointestinal symptoms: The FODMAP approach. J Gastroenterol Hepatol 25:252–258.

19. Gurry T, Nguyen LTT, Yu X, Alm EJ. 2020. Functional heterogeneity in the fermentation capabilities of the healthy human gut microbiota. bioRxiv 2020.01.17.910638.

20. Roels JA. 1983. Energetics and kinetics in biotechnology. Elsevier Biomedical Press.

21. Sawyer CN, McCarty PL, Parkin GF. 2003. Chemistry for Environmental Engineering and Science. McGraw-Hill Education.

22. Gagen EJ, Padmanabha J, Denman SE, McSweeney CS. 2015. Hydrogenotrophic culture enrichment reveals rumen Lachnospiraceae and Ruminococcaceae acetogens and hydrogen-responsive Bacteroidetes from pasture-fed cattle. FEMS Microbiol Lett 362.

23. Rajilić-Stojanović M, de Vos WM. 2014. The first 1000 cultured species of the human gastrointestinal microbiota. FEMS Microbiol Rev 38:996–1047.

24. Owen RW, Spiegelhalder B, Bartsch H. 2000. Generation of reactive oxygen species by the faecal matrix. Gut 46:225–232.

25. Borchers HW. 2019. pracma: Practical Numerical Math Functions.

26. Meeden CJG and GD, Fukuda incorporates code from cddlib (ver 0 94f) written by K. 2019. rcdd: Computational Geometry.

27. Paradis E, Schliep K. 2019. ape 5.0: an environment for modern phylogenetics and evolutionary analyses in R. Bioinformatics 35:526–528.

28. Liaw A, Wiener M. 2002. Classification and Regression by randomForest 2:5.

29. pROC: an open-source package for R and S+ to analyze and compare ROC curves | BMC Bioinformatics | Full Text.

30. Friedman J, Hastie T, Tibshirani R. 2010. Regularization Paths for Generalized Linear Models via Coordinate Descent. J Stat Softw 33:1–22.

31. Preheim SP, Perrotta AR, Martin-Platero AM, Gupta A, Alm EJ. 2013. Distribution-Based Clustering: Using Ecology To Refine the Operational Taxonomic Unit. Appl Environ Microbiol 79:6593–6603.

32. Yu X, Polz MF, Alm EJ. 2019. Interactions in self-assembled microbial communities saturate with diversity. ISME J 1.

33. Søndergaard D, Pedersen CNS, Greening C. 2016. HydDB: A web tool for hydrogenase classification and analysis. Sci Rep 6:34212.

34. Michal B, Gagat P, Jabłoński S, Chilimoniuk J, Gaworski M, Mackiewicz P, Marcin Ł. 2018. PhyMet2: a database and toolkit for phylogenetic and metabolic analyses of methanogens. Environ Microbiol Rep 10:378–382.

35. Zhang H, Yohe T, Huang L, Entwistle S, Wu P, Yang Z, Busk PK, Xu Y, Yin Y. 2018. dbCAN2: a meta server for automated carbohydrate-active enzyme annotation. Nucleic Acids Res 46:W95–W101.

36. Florin T, Neale G, Gibson GR, Christl SU, Cummings JH. 1991. Metabolism of dietary sulphate: absorption and excretion in humans. Gut 32:766–773.

